# Generative AI divide: How College Students’ Backgrounds Affect Their Gen AI Literacy

**DOI:** 10.1101/2024.12.01.626259

**Authors:** Jingbo Jiang, Aiqun Shao

**Author notes:** Corresponding Author:Jingbo Jiang(1997-), male,from Nanping, Fujian, studying for master’s degree in the College of Educational Science, Sichuan Normal University., Aiqun Shao (1974-), Female, from Bazhong, Sichuan, Associate Researcher, Master’s Degree Tutor, College of Educational Science, Sichuan Normal University.

## Abstract

In the field of education, generative artificial intelligence has a profound impact, mainly reflected in areas such as personalized learning, automated assignment evaluation, content generation, and open educational resources. However, some scholars have begun to discuss whether this technology may lead to new inequalities. Therefore, this study aims to explore how user background affects the performance of generative artificial intelligence technology through empirical research while focusing on the potential educational inequalities that may arise in the era of generative AI. The research found that generative artificial intelligence literacy is correlated with grade level, institutional tier, and academic discipline, with higher-grade students, those from prestigious universities, and those studying Applied Sciences and Natural Sciences demonstrating better performance.

## Introduction

Generative AI refers to an artificial intelligence technology that is capable of generating new content based on input data. This technique utilizes deep learning models,particularly Generative Adversarial Networks (GANs),Variational Autocoders (VAEs),and Large Language Models (LLMs),and can be used to generate various types of data such as text,images,audio,video,and more(Cazzaniga et al.,2024). There are several main types of generative AI,the most prominent of which are text generation,image generation,video generation and audio generation(Strobel et al.,2024). Text generation AIs,such as OpenAI’s ChatGPT and Google’s BERT,focus on natural language processing and can generate articles,dialogues or answer questions(Illia and Zyglidopoulos,2023).For image generation,representative tools include DALL-E and Midjourney,which create high-quality images based on text descriptions(Illia et al.,2023).Video generation AI has also grown significantly in recent years,enabling the generation of dynamic video content based on textual descriptions or other inputs. Representative tools include OpenAI’s DALL-E 2 (some of whose features can also be extended to video) and,more recently,Runway ML,which allows users to generate videos with virtual characters or anchors from input text(Elasri et al.,2023). Finally,audio generative AI,such as OpenAI’s Jukedeck and Google’s WaveNet,are capable of synthesizing music or imitating the human voice. These generative AIs show great potential in areas such as creation,design and entertainment.

The impact of generative AI in education is far-reaching,mainly in the areas of personalized learning,automated assignment evaluation,content generation,virtual tutoring,enhanced engagement,open educational resources and teacher training.As a result,some scholars have begun to talk about whether the AI divide will create new inequalities(Capraro et al.,2024;Wilmers,2024).Therefore,with the help of an empirical research paradigm,this paper aims to explore the differences in generative AI literacy among university student groups, and to help students from different subject,educational levels, and institutional backgrounds obtain equal learning opportunities by optimizing the distribution and access of resources. This approach seeks to reduce the educational inequalities arising from generative AI and to build a more inclusive and sustainable education system,ensuring that all students can obtain the necessary educational support in the digital era,thereby contributing to the achievement of sustainable development goals.

### Literature review

#### Digital Divide among Students

The Digital Divide refers to the gap that exists between different social groups in terms of access to and use of information and communication technologies (ICT)(Cullen,2001). This gap is not only in terms of access,such as the lack of Internet connectivity and related equipment in certain regions or populations,but also at the skills level,where some people lack the necessary digital literacy and ability to apply technology(Van and Jan,2006;Maji and Laha,2022).In addition,even with equipment and skills,there may be differences in the quality and diversity of content of the information that people have access to,thus affecting their ability to access information and make decisions (Chowdhury,2004). This phenomenon can exacerbate social inequalities and limit access to education and economic development for certain groups,and thus bridging the digital divide has become an important objective of public policy and social development.

The digital divide also exists among students,and it can affect their academic performance to varying degrees. For instance,computer knowledge and the level of focus on using computers for academic-related tasks have a moderate impact on college students’ learning(Tien and Fu,2008).From the existing review articles,we can see that a series of factors,including students’ socio-economic background,academic major,grade level,and whether they have received digital literacy education,can all influence students’ digital literacy,thereby contributing to the digital divide(Clark,and Gorski,2002;Micheli,2015).There is no doubt that it is extremely important to pay attention to the background of students when exploring the digital literacy divide among college students.

#### Generative AI divide:The new digital divide

As we undergo digital transformation,we’ve come to the age of generative artificial intelligence.The rapid development of Artificial Intelligence has not only changed the way we work,but also profoundly affected our daily lives. From intelligent assistants to creative tools,the widespread use of generative AI provides unprecedented convenience and creativity(Wang et al.,2024).However,many people are confused and uneasy in the face of this new technology,especially in terms of skills and knowledge acquisition,leading to the emergence of a new digital divide(Rana and Verhoeven,2023).

A series of new terms such as the Gen AI Divide and Gen AI Literacy have also been coined to describe the changes that these technological changes are causing in our lives. The Gen AI divide refers to the social inequality caused by differences in technological understanding and application capabilities,where some people and groups may face barriers to accessing,using,or creating technologies and content related to generative AI(Wang and Sun,2024).Gen AI literacy,on the other hand,emphasizes the foundational knowledge,skills,and attitudes that individuals need to have in order to effectively utilize these emerging technologies in this new era(Annapureddy et al.,2024).

With the advent of the generative AI era,the generative ai divide has become the continuation and main manifestation of the digital divide.Research on the digital divide has also gradually shifted from “access”,”use”,and “outcomes” of the Internet to “cognition-access-use-assessment” of generative artificial intelligence (Daepp and Counts,2024;Kitsara,2022).

#### Generative AI Divide and Inequalities

The study of generative AI divide and inequalities was originally referred to in the literature as the “digital inequality”. The study of digital inequality is critical because there are many personal,economic,social,and political benefits to users of the Internet in a variety of contexts(Scheerder et al.,2017).It has resulted in unequal access to education,limited economic participation and increased social exclusion,and has exacerbated barriers to access to information,asymmetries in knowledge and imbalances in regional development,affecting the quality of life of individuals and the overall development of society(Coleman and Blumler,2009;Jennings and Zeitner,2003;Kang and Gearhart,2010;Park and Perry,2008;Polat,2005;Purdy,2017;Smith et al.,2009).

The natural language interaction capabilities and diverse applications of generative artificial intelligence have made it easier for users and have contributed to widespread empowerment. However,the issues of digital inequality arising from the monopolization,dissemination,and limitations of digital technology are a reality that cannot be ignored(Peppiatt,2024).On one hand,the capabilities of generative AI,such as human-like natural language interaction,multimodal interaction,and the ability to solve tasks across various scenarios,have reduced cognitive load and lowered the barriers to usage (De Liso,2023). This helps individuals enhance their productivity,tap into vast resources,and overcome cognitive limitations,presenting valuable opportunities to bridge the gaps between different groups(Cazzaniga et al,2024).On the other hand,issues such as unequal allocation of computational resources,data and algorithmic bias,policy variables like access restrictions,and disparities in human-AI collaboration capabilities can widen the gap between individuals in various aspects of interacting with generative artificial intelligence,particularly amplifying the vulnerabilities of digitally disadvantaged groups(Kim,2021).

However,current research tends to focus on the macro impact of Gen AI on society,with relatively few studies employing empirical methods. Therefore,Therefore,the study aims to explore the following questions.

1. How is the performance of Chinese college students in the use of generative AI literacy?
2. Are there differences in generative AI literacy among Chinese college students from different subject,grades,and institutions?
3. What impact do major,grade,and institution have on generative AI literacy?

### Research design

#### Participants

A total of 530 current college students served as participants,and categorized according to the purpose of the study.In the classification of the institutional level,it is mainly divided into three categories: (1)key universities;(2)General academic universities;(3) Higher vocational colleges.In categorizing the professions,in order to reduce the cumbersomeness of statistics,the researcher referred to other scholars who classified the numerous professions into four main categories(Turner and Bowen,1999):(1)Humanities:Literature,History,Philosophy,Arts;(2)Applied Sciences: Agriculture,Medicine,Engineering;(3)Social Sciences: Economics, Education,Management,Sociology,Law;(4)Natural Sciences:Physics, Chemistry,Biology,Earth and Environmental Sciences.The grade will be recorded according to the participant’s educational background. for example,a second-year undergraduate will be classified as a sophomore,while a third-year master’s student will be classified as a seventh-year student.At the end of the study,the researchers selected 30 participants for follow-up interviews to gain a deeper understanding. The interviews primarily focused on the participants’ development of generative AI usage literacy and how their personal backgrounds influenced this process.

#### Ethical considerations

In terms of research ethics, the researcher followed the principle of informed consent to ensure that participants were fully aware of the purpose, content, and possible impact of the study and voluntarily decided whether or not to participate in the study before presenting the material fragments in the text.

The data does not involve sensitive personal information or commercial interests, and external parties cannot identify personal information from the data, nor does it involve harm to individuals.

#### Questionnaire design and pre-survey

To better assess participants’ skills in using generative AI,the researchers drew upon the EU’s DigComp 2.2 framework and the (Cognition-Access-Utilization-Evaluation framework,hereafter referred to as CAUE framework) generative AI gap identification framework developed by scholar Rong Sun,in order to better explore the differences in skills among users(Sun,2024;Qingyi,2023).

The main changes include the following aspects: (1)The main changes are as follows: The “Information and Data Literacy” domain from the DigComp 2.2 framework has been merged and streamlined with the “cognitive” domain from the CAUE framework into “Tool Cognition Literacy”.(2)Due to the fact that AI usage literacy does not directly involve the following dimensions: communication and cooperation with others in online communities,netiquette,and digital identity,the researchers decided to remove the “Communication and Collaboration” domain from the DigComp 2.2 framework. They incorporated certain dimensions into other domains,such as the ability to distinguish whether one is conversing with a human or a chatbot added to “Tool Cognition Literacy”,and the awareness that everything shared publicly online can be used to train AI systems added to “Evaluation of Results Literacy”.(3)The access domain of the CAUE framework has been merged with certain dimensions from the problem-solving domain of the DigComp 2.2 framework to form “Tool Selection Literacy,” which includes “Identifying Needs and Technological Responses” literacy,”Understanding Cost of Use” literacy,and “Overcoming Access Barriers” literacy.(4)The “Utilization” domain of the CAUE framework has been merged with certain dimensions from the “Digital Content Creation” domain of the DigComp 2.2 framework to form “Content Creation Literacy.”(5)The “Evaluation” domain of the CAUE framework has been merged with the “Safety” domain of the DigComp 2.2 framework to form “Evaluation of Results Literacy”.The final results are presented as follows.The scales all use a 5-point Likert scale,measured from “strongly disagree” to “strongly agree,” with the values increasing sequentially.

In the preliminary research phase,the researcher selected 85 enrolled university students to participate in the pre-study,requiring this group of participants to have high levels of enthusiasm and critical awareness. Some of them later served as research assistants in the subsequent study to help the researcher distribute the questionnaires. The researcher collected 84 survey results through the pre-study,using this data to analyze the reliability and validity of the questionnaire.The Cronbach’s α value of the scale is 0.839,indicating good reliability. The Kaiser–Meyer–Olkin (KMO) coefficient for the entire table is 0.812,which suggests good construct validity of the scale. Furthermore,the results are all significant at the 0.001 level,indicating that the scale has good validity.Finally,the researcher conducted follow-up visits with the participants,and after incorporating their feedback,the questionnaire was collaboratively refined with the assistance of generative AI open-source engineers and doctoral advisors in educational technology, the results are summarized in Table 1.

**Table 1.**
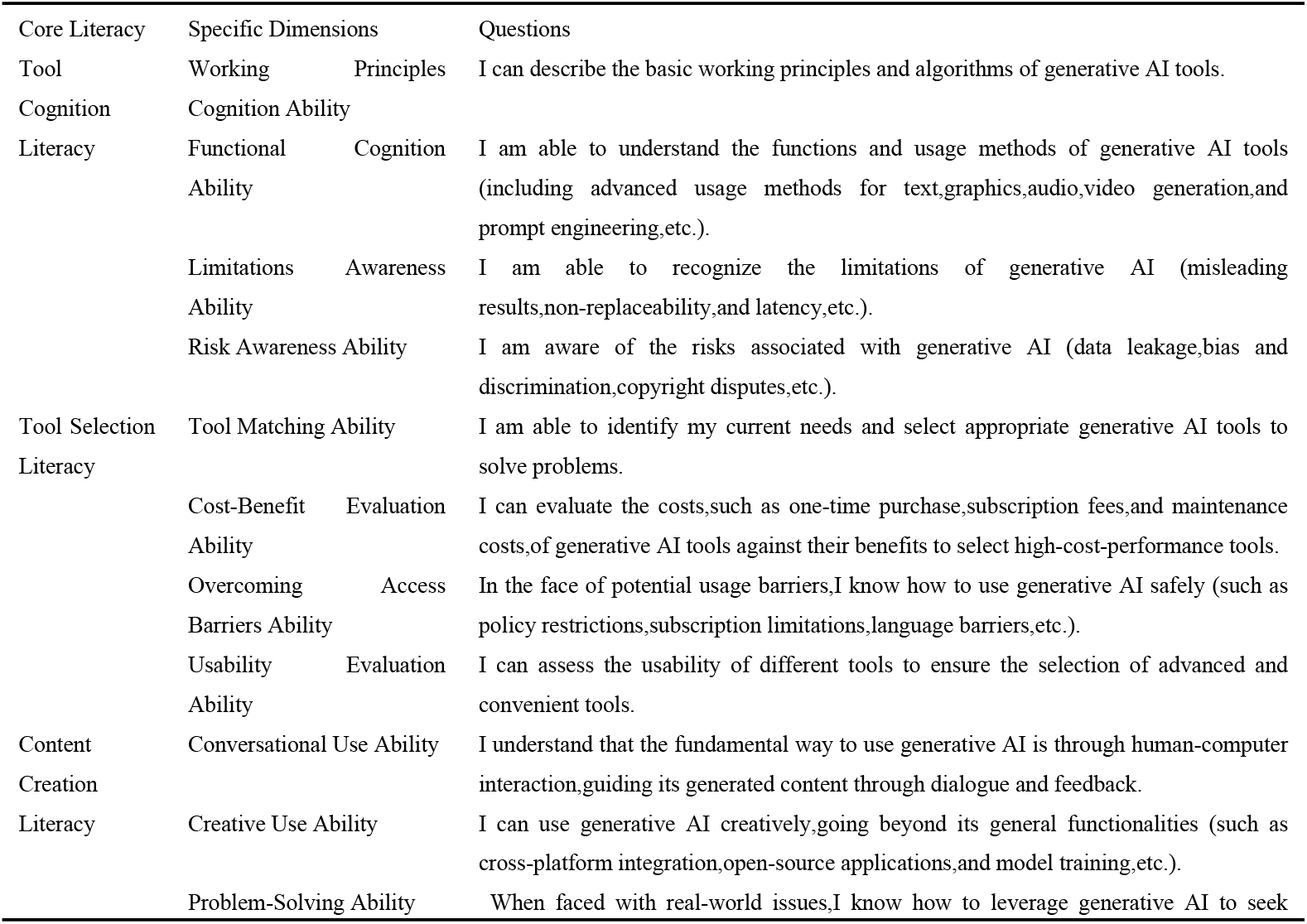
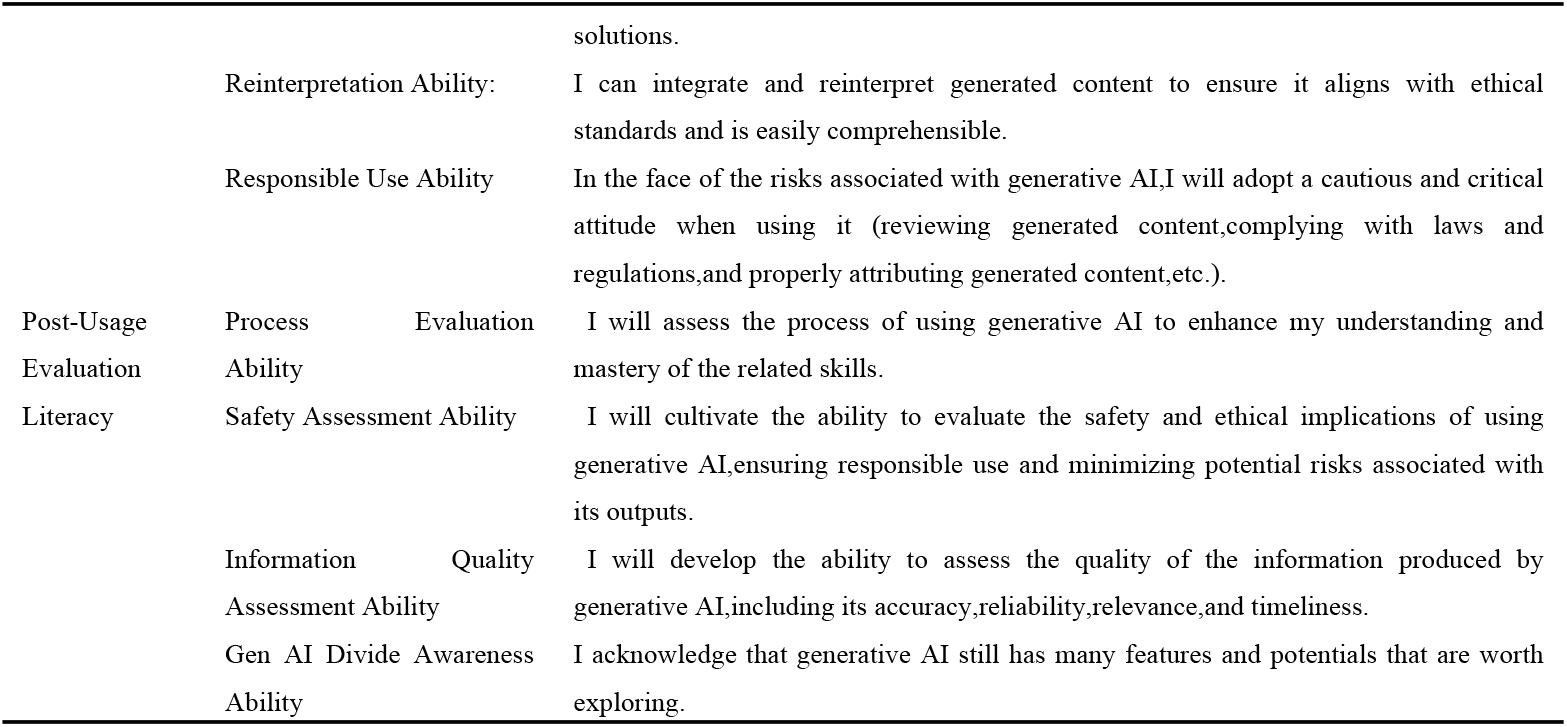
Survey Questionnaire on Generative AI Literacy.

#### Othere Variable Settings

The core variables of the study include three factors: subject,institutional level,and grade. The institutional level is coded from 1 to 3,while the grade variable is encoded from 1 to 12. The subject variable is a categorical variable,and the researcher performed dummy variable encoding for this variable during the analysis. Additionally,considering that previous research has shown that gender,parental occupational status,and urban-rural differences are related to digital literacy to varying degrees,the researcher has set these as control variables. parental occupational status is primarily measured using the Goldthorpe class schema(Chan and Goldthorpe,2007),which is combined into five categories: managers and professionals (Classes I,II,and IVa in the EGP classification),general office staff (Classes IIIa and IIIb),general skilled workers and supervisors (Classes IVb,V,and VI),manual laborers (Class VIIa),and agricultural laborers (Classes IVc and VIIb),coded sequentially from 1 to 5.”The place of residence before the age of 18 is a categorical variable,with urban as 1 and rural as 0. Gender is a categorical variable,with male as 1 and female as 0.”

#### Multicollinearity Test

Since there are no master’s and doctoral students in vocational colleges,and key universities may have a greater number of master’s and doctoral students,there may be a collinearity relationship between grade and institution. Therefore,the researchers conducted a collinearity test on the independent variables.Using SPSS 26.0,it was found that the variance inflation factor (VIF) for the independent variables ranged from 1.255 to 1.476,which is well below the critical value of 5. Therefore,this study model does not have multicollinearity issues.

## Result

### Correlation Analysis

Before conducting regression analysis,in order to better understand the relationships between different variables,the researcher performed a between-subjects effect test. The results are summarized in Table 2.

**Table 2.** Results of the Correlation Analysis.

| Category | F | p |
| --- | --- | --- |
| Grade | 22.835 | 0.000 |
| Subject | 47.839 | 0.000 |
| Institutional Level | 156.318 | 0.000 |
| Grade*Subject | 0.773 | 0.777 |
| Grade*Institutional Level | 0.992 | 0.46 |
| Subject*Institutional Level | 3.187 | 0.041 |
| Grade*Subject*Institutional Level | 0.834 | 0.694 |
| Parental Occupational Status | 2.520 | 0.116 |
| Residence | 6.452 | 0.013 |
| Gender | 18.370 | 0.000 |

The results of the correlation analysis indicate that grade (F=22.835,p < 0.001),subject (F=47.839,p < 0.001),and institution (F=156.318,p < 0.001) have significant effects on the total score. The significance results suggest that grade,subject,and institution all have significant impacts. In terms of interaction effects,only the interaction between subject and institution is significant,while the interactions between grade and subject,grade and institution,and the three-way interaction are not significant.R^2^ = 0.942 (adjusted R^2^ = 0.889).This indicates that the model explains approximately 94.2% of the variance in the dependent variable (total score),and the adjusted R^2^ of 88.9% suggests that the model has a good explanatory power and fits the data well.

### Regression Analysis

Linear regression analysis(Stepwise) was used to examine how participants’ internal variables (background) predict their generative AI literacy,no coefficients were excluded during the regression process,The regression results are summarized in Table 3.

**Table 3.** Results of the Correlation Analysis.

| Category | B | Beta | t | p | Tolerance | VIF |
| --- | --- | --- | --- | --- | --- | --- |
| Grade | 1.456 | 0.450 | 14.096 | 0.000 | 0.693 | 1.442 |
| Institutional Level | 4.973 | 0.444 | 13.692 | 0.000 | 0.670 | 1.492 |
| Humanities | -3.531 | -0.174 | -5.346 | 0.000 | 0.670 | 1.492 |
| Applied Sciences | 1.835 | 0.092 | 2.818 | 0.005 | 0.669 | 1.495 |
| Natural Sciences | 3.607 | 0.175 | 5.350 | 0.000 | 0.664 | 1.507 |
| Residence | 2.457 | 0.136 | 4.347 | 0.000 | 0.720 | 1.398 |
| Gender | 1.839 | 0.105 | 3.749 | 0.000 | 0.908 | 1.101 |
| Parental Occupational Status | 0.359 | 0.054 | 1.814 | 0.071 | 0.791 | 1.161 |

*\*After dummy coding the disciplines, Social Sciences is used as the reference variable.*

The grade variable has a significant positive effect on generative AI literacy,with an expected score increase of approximately 1.456 for each additional grade. The standardized coefficients indicate that the impact of grade on the total score is strong. Institutional factors also have a significant positive effect on generative AI literacy. Differences between institutions can lead to an expected score increase of about 4.973,and the standardized coefficients indicate that its influence is secondary to that of grade.

The unstandardized coefficient (B) for the subject Humanities is −3.531,indicating that compared to the reference variable Social Sciences,participants in the subject Humanities are expected to score 3.531 points lower.The unstandardized coefficient (B) for the subject Applied Sciences is 1.835,indicating that compared to the reference variable Social Sciences,participants in the subject Applied Sciences are expected to score 1.835 points higher.The unstandardized coefficient (B) for the subject Natural Sciences is 3.607,indicating that compared to the reference variable Social Sciences,participants in the subject Natural Sciences are expected to score 3.607 points higher.

### Qualitative Analysis

In selecting the interview subjects,the researchers primarily chose to interview high-scoring participants and low-scoring participants.They also included heterogeneous samples to better understand the situations of those who achieved high scores despite being in less advantageous positions in terms of their majors,institutional levels,and subject,as well as those who,despite being in favorable conditions,obtained low scores.After conducting open coding,axial coding,and selective coding,four categories were ultimately established,as presented in Table 4.

**Table 4.** Results of the Qualitative Analysis.

| Category | Source |
| --- | --- |
| Supporting Factors | Early exposure to digital devices(12), Supportive family environment(5), Strong problem-solving skills(7), Strong self-learning ability(4), Awareness of technological iteration(14), Strong self-efficacy(6), Subject Knowledge Support(15), Equipment Support(6), Peer Support(15), Teacher Support(10), Academic Creation Needs(14), Academic Community Support(8), Personalized Learning Needs(11), Support from experience in using generative AI(16) |
| Obstructive Factors | Long-term residence in rural areas(11), Late exposure to digital devices(8), Poor family economic status(6), Parents opposing the use of electronic devices(7), Lack of personal interest(5), Insufficient self-learning ability(7), Schools maintain a conservative attitude(5), Late exposure to generative AI(12), Difficulty in understanding the principles(5), Simple usage demands(12), Lack of equipment and access tools(8), Lack of guidance(11) |

From the qualitative analysis results,the relationship between generative AI literacy and the level of institutions can be understood from several aspects. Firstly,students at elite universities typically possess better family capital,which means they are exposed to digital devices earlier and can receive family support in the development of their digital skills. Secondly,elite universities have a more open attitude toward generative AI;they allow students to use generative AI through guided applications and provide corresponding assistance. Additionally,students at elite universities often demonstrate stronger self-learning abilities,awareness of technological iteration,and high self-efficacy,which makes them more proactive in engaging with generative AI. Moreover,many respondents mentioned Peer Support,Teacher Support,and Academic Community Support during their process of learning generative AI,which enables students at elite universities to quickly acquire relevant skills even when faced with other challenging conditions by relying on external help.

The relationship between academic disciplines and the use of generative AI can be understood from several aspects. In Applied Sciences and Natural Sciences,many programs are related to computer science or include computer programming within their curricula. This enables students to leverage their disciplinary knowledge to better understand the algorithms,open-source tools,applications,and evaluations associated with generative AI.On the other hand,the concept of personalized learning was mentioned 11 times,with 8 of those mentions coming from students in Applied Sciences and Natural Sciences. In response to the researcher’s follow-up questions,they all indicated that their inclination towards personalized learning might stem from the difficulty of the course material and the strong personalized learning support provided by generative AI. This use of personalized learning has undoubtedly contributed to an increase in these students’ generative AI literacy.

The relationship between grade level and the use of generative AI is easier to understand. On one hand,since the inception of generative AI,there has been a substantial amount of discussion regarding its functions in academic creation. As university students progress in their studies,their needs for academic creation also increasingly rise. This leads to upperclassmen often possessing a higher level of generative AI literacy.On the other hand,as students progress through their grade levels,their practical experience accumulates,allowing them to further develop their generative AI literacy.

## Conclusion

At the dawn of generative AI,many scholars have explored its potential to promote educational equality as well as the risks of increasing inequality (Vázquez-Madrigal et al.,2024;Nedungadi et al.,2024).The analysis of data collected through questionnaires confirmed that the proficiency in using generative AI is related to students’ subjects,grades,and institutional levels,further substantiating the aforementioned viewpoint.The differences in generative AI literacy not only bring varying levels of convenience to the users but also result in differing risks that users face.

On one hand,the application of generative AI fundamentally changes the pattern of knowledge generation.The knowledge generation methods based on big data and algorithms in generative AI can construct knowledge graphs,collect and analyze learning data,track and evaluate the learning process,and identify and diagnose learning problems,thereby providing learners with personalized learning solutions.

On the other hand,the disparities in generative AI literacy have posed numerous risks for marginalized students.A lack of understanding of the working mechanisms and functions of generative AI can lead to insufficient awareness of its limitations and risks,increasing a series of risks such as inefficient use,information silos,dissemination of false information,and privacy leaks.Differences in Tool Selection Literacy for generative AI can make it difficult for users to acquire,evaluate,and match highly usable generative AI tools. This can lead to users falling into information silos and inefficient usage due to inappropriate tools,ultimately impacting their problem-solving capabilities.As the most fundamental literacy, differences in Content Creation Literacy can result in some users demonstrating inadequacies in dialogue, creative use, problem-solving, re-interpretation, and compliance. Almost all risks are associated with this aspect.The lack of Post-Usage Evaluation Literacy may hinder some users from effectively assessing the usage process, information quality, and security after use. This makes it more difficult for them to recognize the potential of generative AI, their own learning directions, and the harms associated with the generative AI divide. Consequently, this can lead to a series of risks, including inefficient usage, information leakage, misleading decisions, intellectual property infringement, continuation of social bias, and dissemination of incorrect information.

**Fig. 1.**
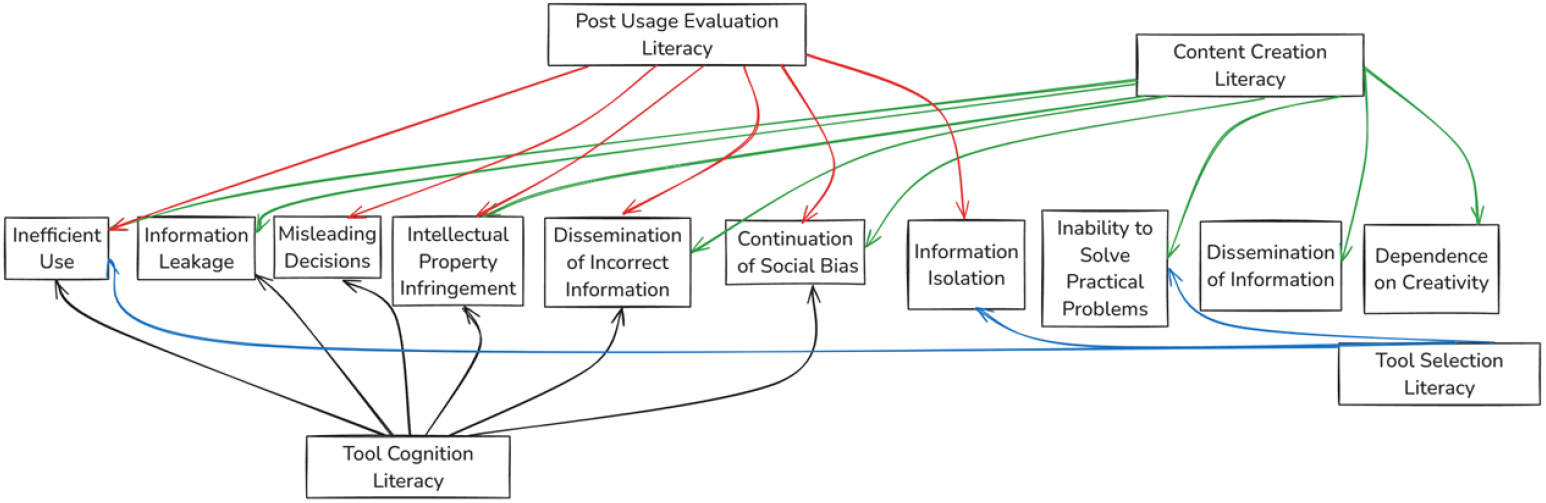
The usage process of generative AI and its associated risks.

In summary, generative AI literacy is correlated with grade level, institutional tier, and academic discipline. Higher-grade students, those from prestigious universities, and those studying Applied Sciences and Natural Sciences performed better in this survey. Therefore, higher education institutions should implement tiered generative AI education tailored to different grade levels, increasing relevant courses and practical opportunities for lower-grade students to further enhance their digital literacy. It is essential to encourage resource sharing and exchange of experiences between different institutions, particularly regarding the development and implementation of generative AI courses, to bridge the human and material resource gaps between institutions and improve overall literacy levels. Additionally, tailored learning plans and practical opportunities should be developed for students in the humanities to enhance their generative AI literacy and reduce the gap with other disciplines. Furthermore, promoting collaboration between faculty and students in Applied Sciences and Natural Sciences with those in the humanities can explore the applications of generative AI across different disciplines and facilitate the intersection and fusion of knowledge.

## Author Contributions

For research articles with several authors, a short paragraph specifying their individual contributions must be provided. The following statements should be used “Conceptualization, JingboJiang and Aiqun Shao.; methodology, JingboJiang; software, JingboJiang; validation, JingboJiang; formal analysis,JingboJiang.; investigation,Aiqun Shao; resources,JingboJiang; data curation,JingboJiang.; writing—original draft preparation, JingboJiang writing—review and editing, JingboJiang;

### Funding

This research was funded by National Education Science “13th Five-Year Plan” 2020 Western Project “Research on Community Education Models for Urban Disadvantaged Groups Based on Community Governance” (Project No.: XKA200291, Host: Shao Aiqun)

## Acknowledgments

In this section, you can acknowledge any support given which is not covered by the author contribution or funding sections. This may include administrative and technical support, or donations in kind (e.g., materials used for experiments).

